# Analysis of coding gene expression from small RNA sequencing

**DOI:** 10.1101/2024.06.21.600062

**Authors:** Aygun Azadova, Anthonia Ekperuoh, Greg N. Brooke, Antonio Marco

## Abstract

The popularity of microRNA expression analyses is reflected by the existence of thousands of sRNA-seq studies where matched total RNA-seq data are often unavailable. The lack of paired sequencing experiments limits the analysis of microRNA-gene regulatory networks. We explore whether protein-coding gene expression can be quantified directly from transcript fragments present in sRNA-seq experiments. We analyze studies containing matched total RNA and small RNA from four human tissues and recover transcript fragments from the sRNA-seq datasets. We find that the expression levels of protein-coding gene transcripts derived from sRNA-seq datasets are comparable to those from total RNA-seq experiments (R^2^ ranging from 0.33 to 0.76). Analyses across multiple tissues and species show similar correlations, indicating that the approach is applicable across organisms. We confirm that transcript half-life and the expression of housekeeping or highly abundant genes do not bias the results. Analysis of the expression of both microRNAs and coding genes from the same sRNA-seq experiments demonstrate that known microRNA-target interactions are, as expected, inversely correlated with the expression profiles of these microRNA-mRNA pairs. For a dual mRNA/miRNA profile, we recommend sequencing the ≥25 nucleotide fraction at ≥ 5 M reads. To confirm the utility of this approach, we apply our method to breast cancer sRNA-seq datasets lacking total RNA-seq data and achieve 75% recall and 64% accuracy comparing inferred coding gene expression with qPCR-validated targets. Our findings demonstrate that quantifying mRNA fragments from sRNA-seq experiments provides a reliable approach to investigate microRNA–mRNA interactions when total RNA-seq is unavailable.

## INTRODUCTION

The analysis of gene expression is a cornerstone of functional genomics. Early works in molecular biology on gene expression were limited, since purified RNA is unstable and difficult to work with. However, the discovery (and use in the laboratory) of reverse transcriptase, permitting the controlled synthesis of RNAs into cDNAs (Maniatis et al. 1976), and the development of microarrays first (Schena et al. 1995), and high-throughput sequencing later (Margulies et al. 2005), boosted our capacity to analyse transcriptomes. Nowadays, the most common technique to analyse gene expression is RNA-sequencing, or RNA-seq. This technique consists of first isolating RNA from a sample, then reverse transcribing into stable cDNA, and finally the sequencing using (mostly) Illumina technology (Bentley et al. 2008). RNA-seq has multiple technical variations, either to identify specific types of transcripts or to characterise other RNA products. For instance, small RNA-sequencing (or sRNA-seq) is a specific technique to sequence small RNAs, mostly microRNAs (Grimson et al. 2007; Ruby et al. 2007). To perform sRNA-seq, a size selection step is introduced, in which cDNA sequences are selected to be in a particular size range. Some sRNA-seq experiments are also coupled with ribosomal RNA (rRNA) depletion to ensure that the samples to be sequenced are enriched in microRNAs and not rRNAs.

MicroRNAs (miRNAs) are short non-coding RNAs approximately 22 nucleotides in length that play a crucial role in post-transcriptional regulation by targeting messenger RNA (mRNA) for degradation or translational repression (Shang et al. 2023). These small but important molecules are involved in a variety of cellular processes, including development, differentiation, and apoptosis (Ratti et al. 2020). The discovery that microRNAs target transcripts by partial pairwise complementarity, permitted the development of multiple target prediction methods (Lewis et al. 2003; Lai et al. 2003; Enright et al. 2003). In mammals, targeted transcripts are in most cases degraded (Baek et al. 2008). Thus, the joint analysis of microRNA and transcript expression can be used to identify microRNA/transcript interactions in combination with other microRNA target prediction programs (van Dongen et al. 2008).

Since the discovery that a deletion of two intronic microRNAs was associated with chronic lymphocytic leukaemia (Calin et al. 2002), the significance of miRNAs in cancer biology has been increasingly recognised (Vannini et al. 2018). The importance of microRNAs in cancer and other diseases is mirrored by the existence of thousands of publications for which sRNA-seq has been performed, either in cells or laboratory-controlled samples, or in patient-derived material. However, in many instances only small RNAs were analysed, and no matched full transcriptome analysis exists. These studies, that utilised precious clinical samples, can be reanalysed as better methods and algorithms are developed to study sRNA-seq. Unfortunately, changes in gene expression associated to changes in microRNA levels cannot be studied in principle as no matching transcriptomic RNA-seq was performed. In this context, we studied whether sRNA-seq experiments contained sufficient fragments from protein-coding transcripts to do a simultaneous analysis of microRNAs and their targets from the same sequencing library. The proposed streamlined approach will not only reduce costs but will also simplify experimental workflows, making the simultaneous analysis of transcriptomes and small RNAs affordable and efficient.

## RESULTS

### Gene expression changes determined using sRNA-seq data correlates with RNA-seq across a range of human tissues

To investigate the potential of sRNA-seq for the analysis of protein coding gene expression, we made use of the sRNA-seq datasets generated by Meunier and collaborators (Meunier et al. 2013). These datasets contain the microRNA expression levels of multiple healthy tissues in several vertebrates. The very same group (Kaessmann laboratory) also generated total RNA-seq datasets from (mostly) the same tissue samples (Brawand et al. 2011). Hence, we first selected four human tissues for which comparable small and total RNA-seq datasets are available: brain, cerebellum, heart and kidney. Both total and small RNA-seq datasets were processed to identify fragments mapping to annotated transcripts. For the analysis we excluded the non-coding RNAs and only considered protein-coding genes. The expression levels between *bona fide* reads from RNA-seq and from fragments derived from sRNA-seq experiments were comparable for all four tissues, especially for brain (Figure 1A), although the association was also comparably high for cerebellum (Figure 1B), heart (Figure 1C) and kidney (Figure 1D). The correlation between gene expression levels from RNA-seq and sRNA-seq libraries was highest in brain (R^2^=0.76) and cerebellum (R^2^=0.62).

**Figure 1.**
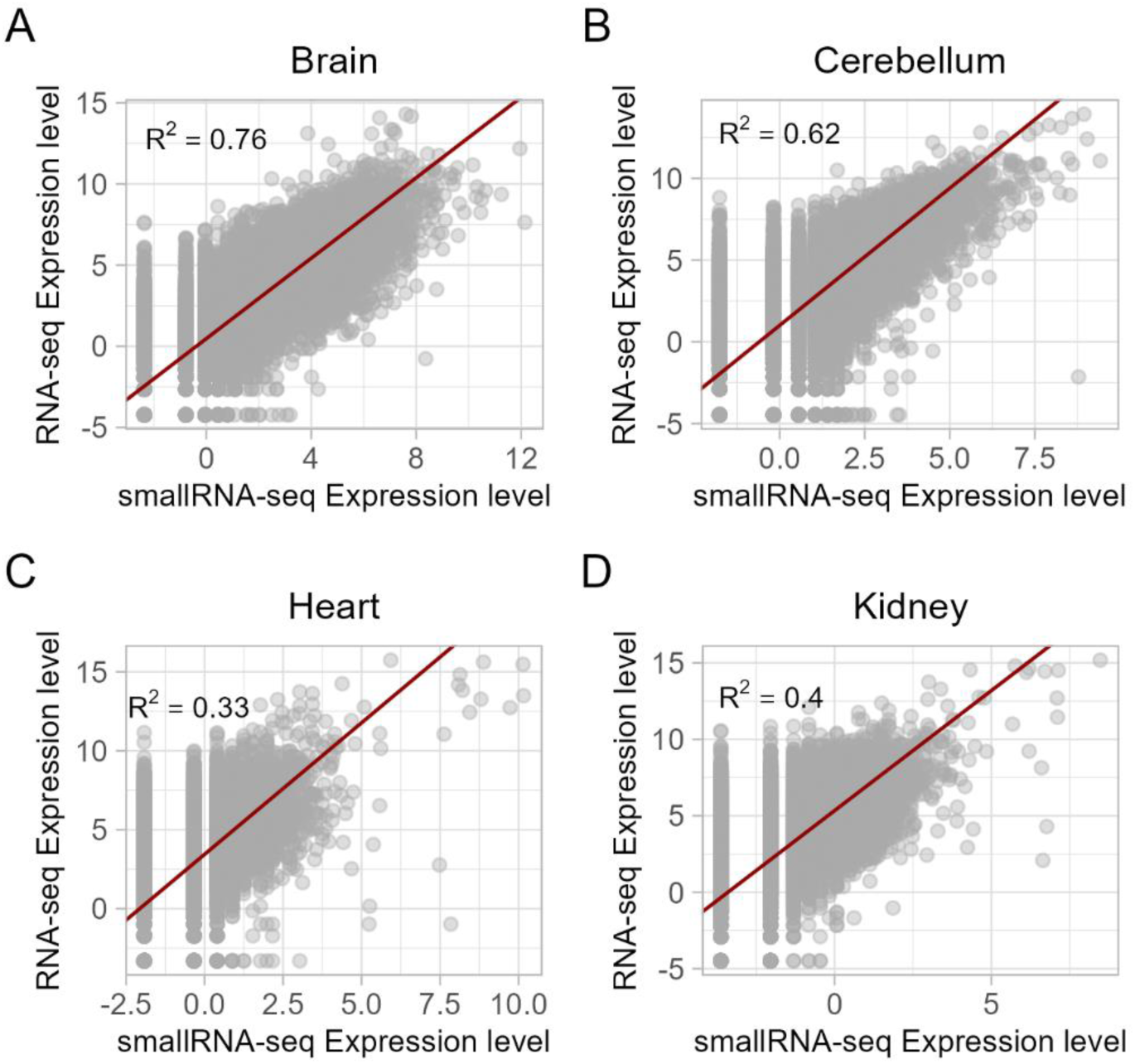
Comparison of the expression levels of protein-coding transcripts from RNA-seq and sRNA-seq matches datasets. Non-coding RNA were removed from the dataset and only protein-coding genes included. The normalised expression level (VOOM transform; see Methods) from matched samples was plotted for brain (A), cerebellum (B), heart (C) and kidney (D) healthy human samples (Brawand et al. 2011; Meunier et al. 2013). A regression line was fitted in all four plots (red line), and the coefficient of determination (R^2^) given within the plot.

### Gene expression changes determined using sRNA-seq data correlates with RNA-seq across a range of tissues from different species

As described above, sRNA-seq can be used to investigate gene expression changes in human tissues. To expand this analysis, we compared the expression level from small RNAs and total RNAs datasets for two additional species: mouse and chicken. For all of the tissues analyzed, the regression fit was significant, and for the majority the coefficient of determination (R^2^) was over 50% (Table 1). More specifically, for heart tissues, in both mice and chickens the association was particularly high (R^2^ approximately 70%).

**Table 1.** Association between gene expression inferred from small RNA datasets and total RNA expression levels. R^2^ were derived from comparing the expression values for coding genes extracted from small (Meunier et al. 2013) and total (Brawand et al. 2011) RNA libraries (Regression test).

| Species | Tissue | $R^2$ | p |
| --- | --- | --- | --- |
| mouse | Brain | 0.517 | <0.001 |
| mouse | Cerebellum | 0.530 | <0.001 |
| mouse | Heart | 0.692 | <0.001 |
| mouse | Kidney | 0.475 | <0.001 |
| mouse | Testis | 0.409 | <0.001 |
| chicken | Brain | 0.561 | <0.001 |
| chicken | Cerebellum | 0.448 | <0.001 |
| chicken | Heart | 0.713 | <0.001 |
| chicken | Testis | 0.292 | <0.001 |

Some samples showed higher R^2^ values than others, and these were different across species. For instance, human brain expression is better captured by sRNA-seq than heart expression, but this pattern is reversed in mice. This may be due to differences in sequencing depth across samples. However, there is no association between sequenced reads per genome megabase and R^2^ (R = -0.016, p = 0.958, Pearson’s correlation test; Supplementary Table 1). However, if we consider only reads that after adapter removal were longer than 25 nucleotides (unlikely to be microRNAs), a clear association between number of sequenced reads and R^2^ becomes clear (R = 0.663, p = 0.026; Supplementary Table 1), if we exclude the two testes samples. This is expected since testes are rich in piRNAs, which are longer than microRNAs and likely to represent a significant fraction of the analysed reads (Sun et al. 2022). In addition, given the potential impact of repeated sequences, we also investigated the relative role of read complexity in the sequencing libraries. To do so, we computed the Normalised Shannon Entropy based in the uniqueness of sequencing reads (ranging from 0 if all reads are the same to 1 if all reads are unique) and built a multivariate linear model to regress R^2^ with two independent variables: complexity and sequencing depth (Supplementary Table 1). When all samples are considered, we found no association between either sequencing depth or complexity and R^2^ (depth: p=0.158; complexity: p=0.587). If the two testes samples are removed (as above), only sequencing depth seems to be associated with R^2^ (complexity: p=0.319; depth: p=0.0.031). From this analysis we conclude that, although sequence complexity may have an impact, sequencing depth is a major determinant of whether small RNA libraries can be used to successfully measure coding-genes expression levels.

### Transcript half-life has a negligible effect on small-RNA expression estimates across tissues

To evaluate potential biases in the transcript coverage between total and small RNA libraries, we computed the relative enrichment of mapped reads in untranslated regions compared to the coding sequence for each transcript. For the four human total RNA sets analysed previously, there was a clear enrichment in reads mapping to the CDS compared to both 3’ UTR and 5’ UTR. In small RNA libraries there were mixed results: for heart and kidney, there was an enrichment in reads mapping to both UTRs, in cerebellum an enrichment in CDS and 3’ UTR compared to 5’ UTR, an in brain the pattern was comparable to total RNA libraries (enrichment in CDSs, Supplementary Figure 1). This result is consistent with the better coverage of non-microRNA sequences in small RNAs from brain compared to other tissues.

The expression levels quantified from the small RNA libraries could be associated with degradation fragments from transcripts. We explored this by quantifying the impact of total RNA levels (RNA-seq expression) and the half-life of transcripts in the estimation of gene expression from small RNA libraries. For brain samples, there is a significant inverse association between half-life and the gene expression levels quantified from sRNA sequencing (p=0.000017; Supplementary Table 2) but the effect size is negligible (slope = -0.017) compared with the impact of the total RNA level of the gene transcript (p<0.00001, slope = 0.659). The interaction term of the regression is negligible. A comparable effect is observed in the other studied samples (Supplementary Table 2). In conclusion, although the half-life of transcripts does have a significant effect on the estimated expression values from small RNA libraries, the size effect is negligible compared to the actual expression levels from total RNA libraries. Transcript half-life is therefore unlikely to impact/bias our analysis pipeline.

### The association between the differentially expressed genes identified using sRNA-seq and RNA-seq is not due to co-detection of housekeeping genes

The association between small RNA and total RNA expression could be partly due to the co-detection of highly expressed housekeeping genes. To rule this out, we functionally annotated the top 10% of the highest expressed genes form the small RNA inferences from human samples. Table 2 shows the top 3 most enriched terms for Cellular Component (Gene Ontology) and KEGG pathways. The enriched terms were consistent with the expected functional features of the analysed datasets. For instance, both brain and cerebellum expression from small RNAs were enriched in genes associated with neural cell body or presynapse; cerebellum was specifically enriched in glutamatergic synapse (a type of synapse enriched in cerebellum); and brain was enirhced in thyroid hormone signalling, which has a more important role in adult brain that in cerebellum. Likewise, kidney is enriched in lysine degradation, which predominantly occurs in liver and kidney (Vaz and Wanders 2002) and heart is enriched in hypertrophic cardiomyopathy associated genes (Sorajja et al. 2000).

**Table 2.** CC and KEGG enrichment analysis for highly expressed genes from small RNA datasets. CC (GO): Cellular Component (Gene Ontology).

| **Tissue** | **Annotation** | **Enriched term** | **Enrichment** | **q** |
| --- | --- | --- | --- | --- |
| brain | KEGG | Long-term potentiation | 3.8 | 1.59E-10 |
| brain | KEGG | Thyroid hormone signaling pathway | 2.9 | 9.69E-10 |
| brain | KEGG | Circadian entrainment | 3.1 | 4.10E-09 |
| cerebellum | KEGG | Glutamatergic synapse | 3.5 | 3.50E-14 |
| cerebellum | KEGG | Circadian entrainment | 3.4 | 4.26E-11 |
| cerebellum | KEGG | Long-term potentiation | 3.9 | 7.63E-11 |
| heart | KEGG | Focal adhesion | 3.2 | 1.05E-13 |
| heart | KEGG | Proteoglycans in cancer | 2.9 | 2.85E-10 |
| heart | KEGG | Hypertrophic cardiomyopathy (HCM) | 3.7 | 5.84E-08 |
| kidney | KEGG | Focal adhesion | 2.3 | 1.51E-07 |
| kidney | KEGG | Lysine degradation | 3.6 | 3.35E-07 |
| kidney | KEGG | Adherens junction | 3.3 | 3.35E-07 |
| brain | CC (GO) | neuronal cell body | 2.6 | 0 |
| brain | CC (GO) | presynapse | 2.7 | 0 |
| brain | CC (GO) | cytoplasmic region | 2.6 | 0 |
| cerebellum | CC (GO) | neuronal cell body | 2.8 | 0 |
| cerebellum | CC (GO) | presynapse | 3.0 | 0 |
| cerebellum | CC (GO) | cytoplasmic region | 2.4 | 0 |
| heart | CC (GO) | actin cytoskeleton | 2.9 | 0 |
| heart | CC (GO) | cell-substrate junction | 3.5 | 0 |
| heart | CC (GO) | cell-substrate adherens junction | 3.5 | 0 |
| kidney | CC (GO) | cell-substrate junction | 3.0 | 0 |
| kidney | CC (GO) | cell-substrate adherens junction | 3.0 | 0 |
| kidney | CC (GO) | focal adhesion | 3.1 | 0 |

### MicroRNA expression is associated with target gene expression quantified using sRNA-seq

Given that we successfully measured coding gene expression levels from fragments present in sRNA-seq datasets, we also quantified the expression level of microRNAs and compared the expression profile between all microRNA/coding-gene pairs from the same sRNA-seq. Then, we identified those pairs with known interactions previously described in the literature as compiled in miRTarBase (see Methods). In this analysis, we used all the samples available for small RNAs which include, on top of the four human tissues studied already, a sample from testes. By plotting the ratio of known microRNA-target interactions as a function of the expression correlation, we clearly observe that anti-correlated pairs are enriched in target interactions, while highly correlated pairs show a paucity of targets (Figure 2). This indicates that the joint analysis of microRNAs and their potential targets, solely generated from sRNA-seq experiments, can be used to study the function of microRNAs in specific tissues.

**Figure 2.**
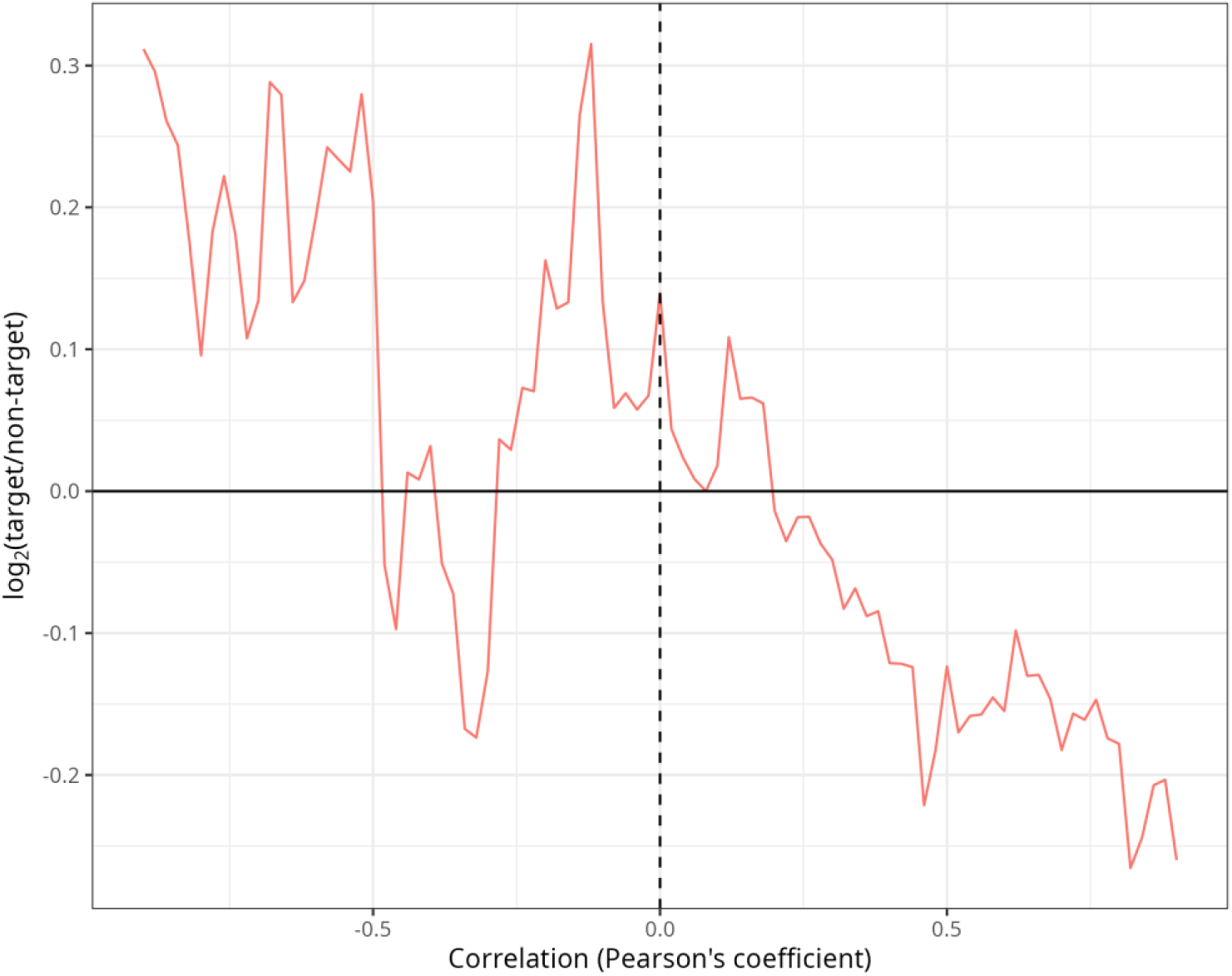
Co-expression of microRNAs and their targets within sRNA-seq experiments. For all pairwise comparisons between a microRNA and a protein-coding gene, the x-axis gives the correlation of expression values in the human samples in Figure 1, and the y-axis shows the associate proportion of validated target pairs with respect to the total number of pairs in the bin. Each value in the x-axis is a bin of size +/- 0.2 around the x-axis value in a sliding window analysis with step size of 0.02.

### Gene expression analysis, performed on small RNA patient datasets, successfully identifies genes linked to breast cancer

To evaluate whether this methodology can extract useful information from clinical samples, we studied 24 samples from 12 patients with breast cancer sequenced by (Meerson et al. 2019); this study focused on microRNAs and only sRNA-seq was performed. This allowed us to perform a paired analysis (two conditions: matched tumour versus non-tumour) to identify genes differentially expressed in breast cancer. The expression levels of microRNAs and coding-genes were quantified from the sRNA-seq datasets, and we performed differential gene expression analysis for both. The analysis of microRNAs reveals that dozens of microRNAs are differentially expressed between tumour and non-tumour samples correcting for batch (patient). More specifically, for a false discovery rate or 1% and a log_2_ fold-change difference of at least 1/-1, we identified 54 upregulated and 29 downregulated microRNAs in breast cancer samples. Among microRNAs, the most significant changes are for *MIR144* (downregulated in tumours) and *MIR429* (upregulated in tumours) (Supplementary Figures 2 and 3).

**Figure 3.**
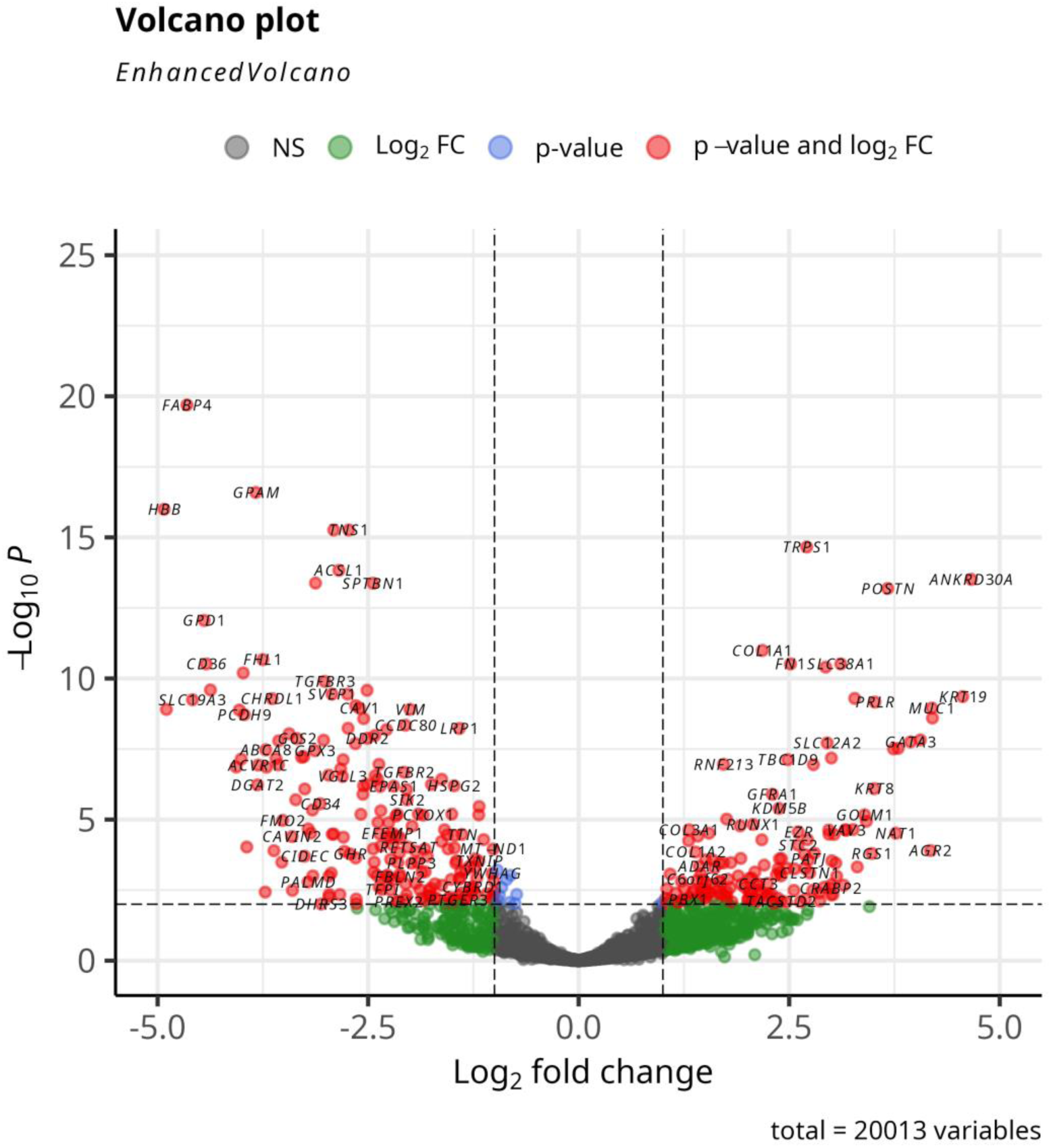
Differential gene expression of protein-coding genes from breast cancer paired sRNA-seq experiments. Volcano plot representing the expression fold change (DESeq2) on the x-axis of paired breast cancer samples (see main text) against the -log_10_ of the q value (FDR corrected p value) generated during the differential gene expression analysis. Identified differentially expressed genes are shown in red.

Importantly, the limited number of reads mapped to coding-genes in the sRNA-seq was sufficient to permit differential gene expression analysis, and many coding-genes were found to be up- and downregulated in tumours (Figure 3 and Supplementary Figure 4). The functional annotation of upregulated genes from this analysis reveals an enrichment in functional categories (within the Biological Process domain in Gene Ontology) related with cell proliferation, as expected for cancer samples (Table 3). Also, the annotation to disease related databases (OMIM and Glad4U) consistently shows an enrichment in breast cancer related categories (Table 4). These results confirm that the differentially regulated genes identified from the sRNA-seq experiments are consistent with those expected from breast cancer samples.

**Table 3.**
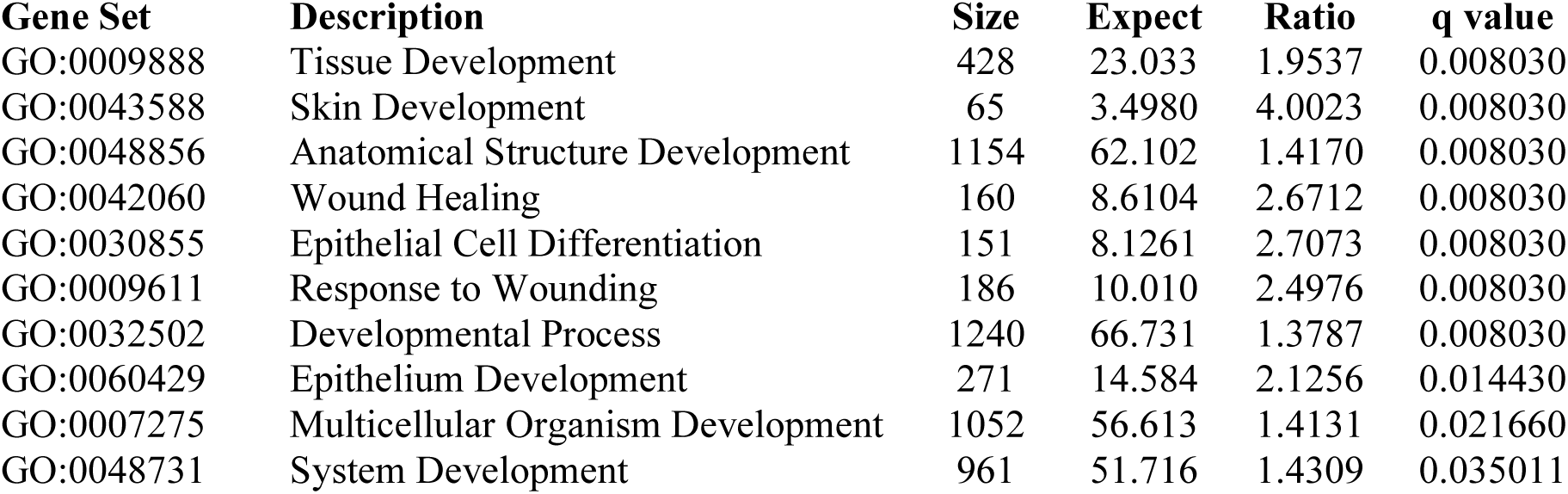
Gene Ontology enrichment analysis for upregulated genes. Size: size of the category; Expect: expected number of genes from query in category; Ratio: ratio of observed over expected number of genes.

| Gene Set | Description | Size | Expect | Ratio | q value |
| --- | --- | --- | --- | --- | --- |
| GO:0009888 | Tissue Development | 428 | 23.033 | 1.9537 | 0.008030 |
| GO:0043588 | Skin Development | 65 | 3.4980 | 4.0023 | 0.008030 |
| GO:0048856 | Anatomical Structure Development | 1154 | 62.102 | 1.4170 | 0.008030 |
| GO:0042060 | Wound Healing | 160 | 8.6104 | 2.6712 | 0.008030 |
| GO:0030855 | Epithelial Cell Differentiation | 151 | 8.1261 | 2.7073 | 0.008030 |
| GO:0009611 | Response to Wounding | 186 | 10.010 | 2.4976 | 0.008030 |
| GO:0032502 | Developmental Process | 1240 | 66.731 | 1.3787 | 0.008030 |
| GO:0060429 | Epithelium Development | 271 | 14.584 | 2.1256 | 0.014430 |
| GO:0007275 | Multicellular Organism Development | 1052 | 56.613 | 1.4131 | 0.021660 |
| GO:0048731 | System Development | 961 | 51.716 | 1.4309 | 0.035011 |

**Table 4.** Disease categories enrichment for upregulated genes in OMIM and GLAD4U. ^1^OMIM; ^2^GLAD4U. Headers as in Table 3.

| Gene Set | Description | Size | Expect | Ratio | q value |
| --- | --- | --- | --- | --- | --- |
| 114480<sup>1</sup> | Breast Cancer | 7 | 0.026 | 76.095 | 0.001574 |
| 176807<sup>1</sup> | Prostate Cancer | 7 | 0.026 | 38.048 | 0.069562 |
| 601626<sup>1</sup> | Leukemia, Acute Myeloid | 7 | 0.026 | 38.048 | 0.069562 |
| PA446482<sup>2</sup> | Skin and Connective Tissue Diseases | 113 | 6.396 | 4.8466 | 1.382e-11 |
| PA445676<sup>2</sup> | Skin Diseases | 121 | 6.849 | 4.5262 | 5.572e-11 |
| PA443560<sup>2</sup> | Breast Neoplasms | 140 | 7.925 | 3.7857 | 1.574e-8 |
| PA443559<sup>2</sup> | Breast Diseases | 128 | 7.245 | 3.7266 | 2.040e-7 |
| PA445062<sup>2</sup> | Neoplasms | 223 | 12.62 | 2.7728 | 1.578e-6 |
| PA447242<sup>2</sup> | Epithelial Cancers | 124 | 7.019 | 3.5618 | 1.813e-6 |
| PA446646<sup>2</sup> | Carcinoma, Ductal, Breast | 34 | 1.925 | 6.7549 | 2.223e-6 |
| PA165108776<sup>2</sup> | Infiltrating Duct Carcinoma of Breast | 34 | 1.925 | 6.7549 | 2.223e-6 |
| PA445058<sup>2</sup> | Neoplasm Metastasis | 157 | 8.887 | 3.1507 | 2.399e-6 |
| PA443610<sup>2</sup> | Carcinoma | 159 | 9.000 | 3.1111 | 2.901e-6 |

### Analysis of gene expression data in sRNA-seq can be used to identify/validate microRNA target regulation

To analyse the potential of our approach for identifying microRNA target sites, we first identified canonical sites in microRNA–transcript pairs. We then quantified enrichment as the proportion observed in experimentally validated targets relative to non-validated targets. More specifically, we computed the log-odds ratio of the proportion of validated target sites for: (i) upregulated and downregulated microRNAs compared to downregulated transcripts; (ii) and upregulated and downregulated microRNAs compared to upregulated transcripts. Our results indicate that for downregulated microRNAs there is a statistical enrichment in validated targets compared to upregulated microRNAs, when we considered downregulated transcripts (p=0.0025; Table 5). This indicates that when we identify pairs of down-miR:up-transcript we can identify functional target sites. However, in the reverse case (up-miR:down-transcript) the association was not statistically significant, although there was an enrichment in targets (Table 5).

**Table 5.**
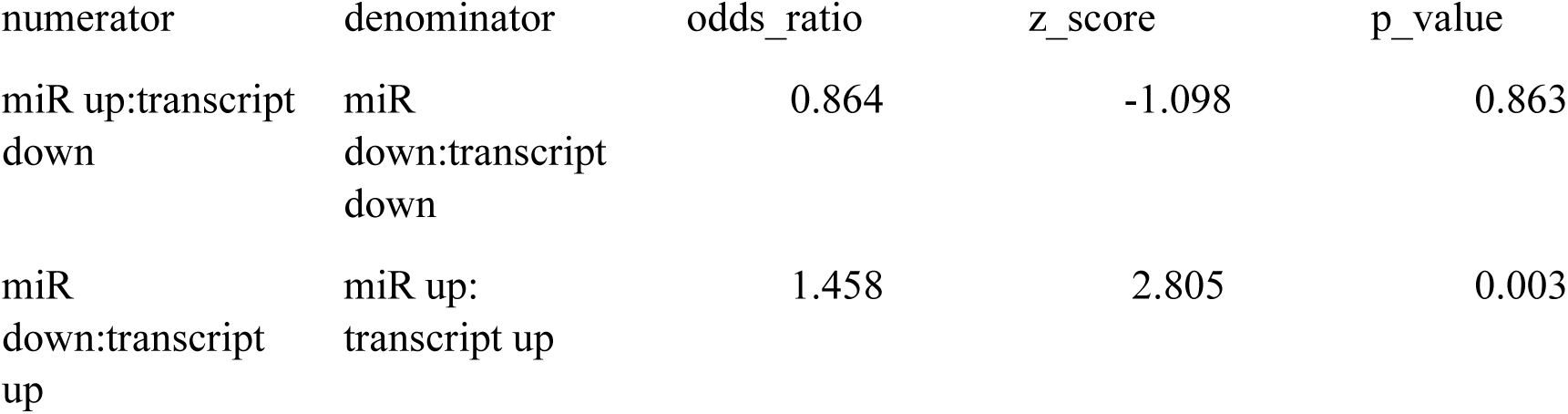
Enrichment in validated targets for differentially expressed microRNAs.

| numerator | denominator | odds_ratio | z_score | p_value |
| --- | --- | --- | --- | --- |
| miR up:transcript<br>down | miR<br>down:transcript<br>down | 0.864 | -1.098 | 0.863 |
| miR<br>down:transcript<br>up | miR up:<br>transcript up | 1.458 | 2.805 | 0.003 |

In the original paper studying small RNAs in breast cancer samples, the authors validated the targets of miR-10b-5p using qPCR. They considered 15 known targets: *BCL2L11, BDNF, CDKN1A, CDKN2A, HOXD10, KLF4, MAPRE1, NCOR2, PAX6, PIEZO1, PPARa, PTEN, SRSF1, TP53* and *TRA2B*. Of those, they found anti-correlated expression of miR-10b-5p with *MAPRE1, PIEZ01, SRSF1* and *TP53*. We used this data to validate our approach by analysing our gene expression calculated levels from small RNA libraries; focussing on genes that have an opposite expression fold change (cancer versus normal) with respect to miR-10b-5p (Figure 4). For all 14 genes (*PAX6* was excluded as we did not detect any reads mapped to it) we found that three out of the four validated targets were upregulated in our analysis (miR-10b-5p is downregulated in breast cancer), representing a 75% recall. Considering all of the 14 predicted targets, we also computed a precision of 43% and an accuracy of 64%. If we increase the log_2_ fold-change threshold to determine which genes are upregulated according to our DGE analysis and compare it again to the gold standard, we observed, as expected, that the recall decreases and the precision increases. However, the accuracy remains high for log_2_ fold-change thresholds between 0 and 2, with values around 70% (Supplementary Figure 5).

**Figure 4.**
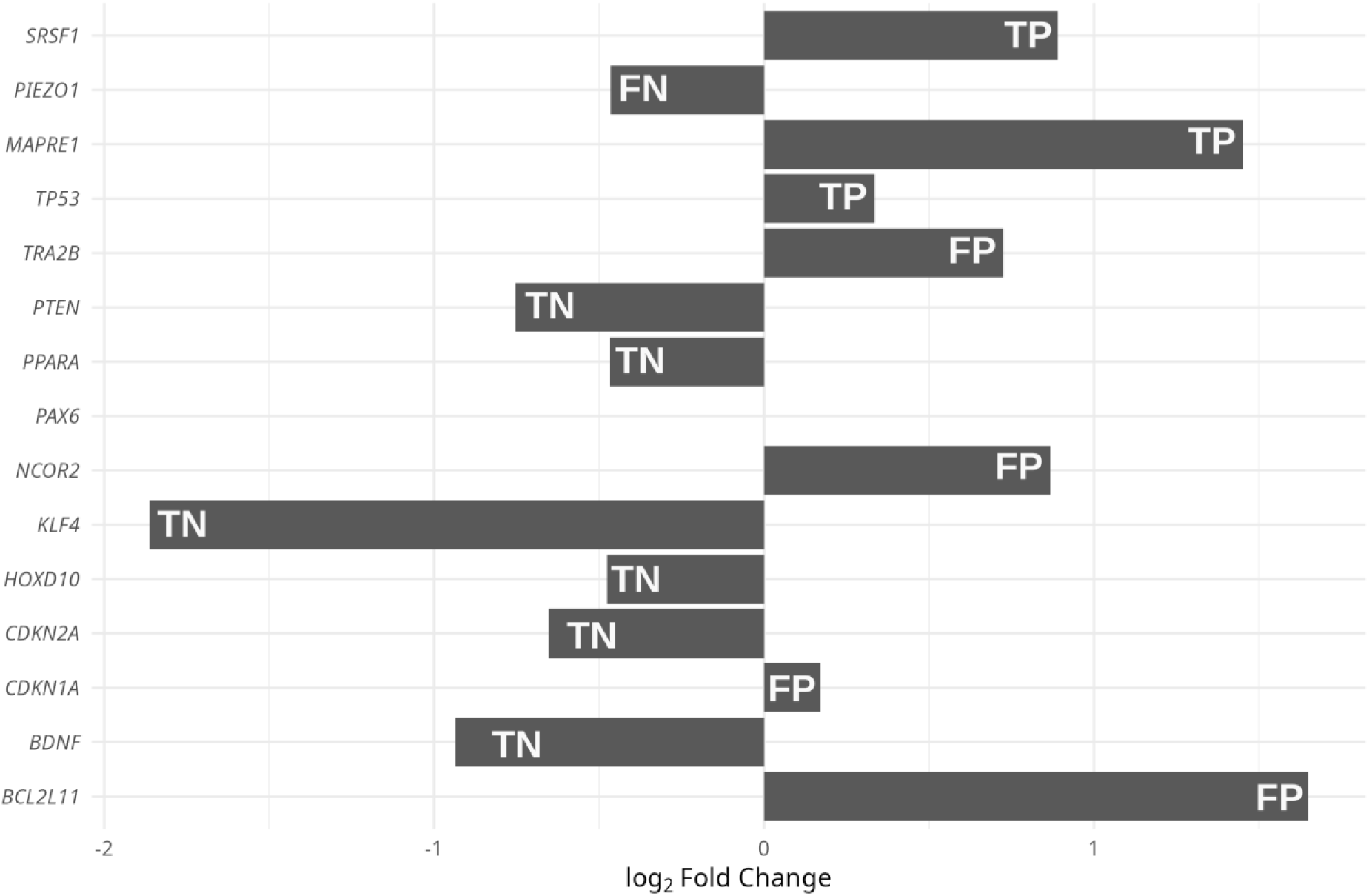
Fold-change of miR-10b-5p target genes. Fold change of target genes inferred from small RNA libraries. The first four (top) genes are validated miR-10-5p targets according to (Meerson et al. 2019). All other are non-validated targets. Genes are labelled, according to how their expression level compared to the expected from the gold-standard, as true positives (TP), false positives (FP), true negatives (TN) and false negatives (FN).

This, together with the previous analysis, suggests that the use of small RNA datasets can be used to identify and validate microRNA targets.

## DISCUSSION

In this work we first compared the expression levels of protein-coding sequences from matched RNA-seq and sRNA-seq experiments across different vertebrate species and in multiple tissues, to investigate if the fragments present in sRNA-seq can be used to recover biologically and clinically meaningful mRNA signatures, therefore enabling dual analysis (miRNA and mRNA) from a single library and sequencing run. Importantly, our approach captures >50% of variance across these different sample types. We also observed good correlations in our analysis of mouse and chicken datasets, confirming that the usefulness of our approach is not limited to human data.

The association between RNA-seq and our mRNA predictions from sRNA-seq was different for different tissues and not always consistent across species. We showed that a high association (determination coefficient) is associated with a high sequencing depth when we exclude reads of size 25nt or smaller (likely microRNAs) except for testis samples (piRNAs). In addition, we showed that read complexity of sRNA-seq experiments is not a determinant either. It is therefore recommended that ≥ 25 nucleotide fraction at a sequencing depth of ≥ 5 M reads should be used per sample when performing sRNA-seq for dual profiling. In contrast, degradation bias (half-life analysis) and noise from housekeeping genes do not appear to have a significant effect upon/bias our approach. For the latter, Gene Ontology analysis identified tissue-specific pathways reinforcing that the signal is biologically specific rather than dominated by ubiquitous transcripts.

We also analysed paired breast cancer and normal samples for which only small RNA sequencing data were available, and we successfully identified differentially expressed microRNAs and coding-genes, as well as identifying potential microRNA-transcript interactions. Our approach was confirmed using microRNA targets that have been previously validated (Meerson et al. 2019). Validation against qPCR-confirmed miR-10b-5p targets achieved 75% recall and 43% precision, illustrating practical utility for target nomination from sRNA-seq alone.

Although the computational prediction of microRNA targets has been relatively successful in the past (see Introduction), experimental techniques have improved our ability to detect/confirm bona fide targets. These techniques are diverse and include variations of immunoprecipitation and expression analysis (reviewed in Thomson et al. 2011). The joint analysis of gene expression of both microRNAs and their potential targets has been successfully used in the past, starting with the pioneering work by Huang et al. (2007). In that article, the authors describe a method that builds regulatory networks by combining the expression profile of matched miRNA/mRNA microarray experiments. Ever since, other studies using RNA-seq/sRNA-seq matched experiments have been used to identify microRNA targets (e.g. Jacobsen et al. 2013). A step further, in part to avoid unwanted effects due to differences in sample/library preparation, is the use of the same high-throughput expression experiment to study simultaneously both microRNAs and their potential targets. Two works from the Banfi laboratory exploited this idea in two different ways (Gennarino et al. 2009, 2012). First, considering that many intronic microRNAs have their expression linked to that of their host gene (Baskerville and Bartel 2005), they use gene expression microarrays to identify genes with anti-correlated expression with the host gene, as a proxy of intronic microRNA/target interaction (Gennarino et al. 2009). Second, they considered, again using microarray experiments, that targets of the same microRNA are co-expressed, and use this to identify microRNA targets (Gennarino et al. 2012). In this work we go a step further and study, as far as we are aware, for the first time, simultaneously the expression level of microRNAs and their potential targets from the same sRNA-seq experiments.

The use of anti-correlation between microRNAs and their specific targets as a means of identifying potential, biologically relevant regulatory interactions, although widely employed (including in the present study), has certain limitations. First, microRNA targeting is a post-transcriptional regulatory mechanism and, when microRNA–target pairing is partially complementary, as is typically the case in animals, protein synthesis is repressed (Bartel 2009). However, target RNA stability is also influenced by microRNA targeting, and, particularly for microRNAs exerting strong regulatory effects, target degradation is expected (Selbach et al. 2008; Baek et al. 2008), thereby justifying the use of transcriptomic approaches to study microRNA targets. Second, the regulatory impact of microRNAs can be complex and involve multiple regulatory steps. In this context, anti-correlation between a microRNA and a transcript does not necessarily indicate the presence of a functional target site. Regulatory network–aware tools, such as those described above (Huang et al. 2007), account for the transcriptional responses of multiple microRNAs and transcripts. Future iterations of our method should therefore incorporate this capability.

Further work is necessary to better understand the factors that will allow for the systematic analysis of both microRNAs and their targets from the same sRNA-seq experiments, but from the outcomes of this work, this approach is valid and will provide useful data to better understand microRNA-target gene regulatory networks. It will also enable users to predict gene expression changes from published datasets where only small RNA-seq data are available, and from future studies in which resources or samples are limited, ensuring that the maximum amount of information is extracted from each experiment.

## METHODS

### Datasets and databases

The total tissue RNA expression datasets were those from (Brawand et al. 2011) available at https://www.ebi.ac.uk/ena with accession PRJNA143627. The small RNA human tissue expression datasets are from (Meunier et al. 2013) with accession number PRJNA174234. The small RNA and total RNA experiments were performed by the same group. Gene annotations (CDS and UTRs) are from Ensembl version 113 (October 2024) retrieved with biomaRt (Kinsella et al. 2011). Breast cancer (patients) data was retrieved from (Meerson et al. 2019) with accession PRJNA494326. Experimentally validated microRNA targets are from miRTarBase [v 9.0] (Chou et al. 2018), and canonical microRNA targets were predicted using seedVicious [v1.3] (Marco 2018).

### Analysis of coding gene and microRNA expression

Adaptors were removed from reads with cutadapt [v3.7] (Martin 2011) and reads were mapped to the human genome hg38 with HISAT2 [v2.2.1] (Kim et al. 2019) with default parameters. We then used featureCounts v2.0.2 (Liao et al. 2014) to count the number of reads in each feature. For human transcripts we used the annotation in GENCODE [v43] (Frankish et al. 2019) and for microRNAs we used miRBase 22.1 (Kozomara et al. 2019). When comparing the expression profile of RNA-seq and sRNA-seq experiments for the same tissues/samples, we first used the Voom transformation on read counts (Law et al. 2014) using limma 3.52.2 (Ritchie et al. 2015). The computation of log_2_ fold-change expression values, and the differential gene expression analysis, were done with DESeq2 (v1.36.0) (Love et al. 2014), including a patient term in the model (*expression ∼ patient + tumour*). For the analysis of transcript half-life, we used the information from Tani et al. (2012), and then build a regression model *small_RNA ∼ total_RNA + half-life + interaction* to evaluate the relative impact of transcript half-life compared to the transcript abundance based on levels from the total RNA libraries. All statistical analyses and figures were done using R 4.2.1 (R Development Core Team 2004). Functional annotation was performed with WebGestalt 2024 (Elizarraras et al. 2024) setting a minimum number of elements per category to 5 and using the list of genes that passed the DESeq2 default filtering as the background list, leaving other options as default. The categories evaluated were Biological Process in Gene Ontology (The Gene Ontology Consortium 2000), OMIM (Hamosh et al. 2005) and GLAD4U (Jourquin et al. 2012). Volcano plots were drawn with the EnhancedVolcano R package (Blighe et al. 2024).

### Software availability

All results generated and the scripts are available from GitHub at https://github.com/antoniomarco/CDS_from_sRNAseq and as a Supplemental Code file.

## COMPETING INTEREST STATEMENT

The authors declare no competing interests.

## ACKNOWLEDGMENTS

The authors acknowledge the use of the High-Performance Computing Facility (Ceres) and its associated support services at the University of Essex in the completion of this work. Author contributions: AM and GNB conceived and devise the project, interpreted the results and wrote the manuscript; AE, AA and AM performed the analyses. Funding: This work was funded by a BBSRC Impact Accelerator Award BB/X511171/1. AA was funded by the Ministry of Science and Education, Republic of Azerbaijan. GNB is also supported by BBSRC grants BB/W020033/1 and BB/X018997/1. For the purpose of open access, the author(s) has applied a Creative Commons Attribution (CC BY) licence to any Author Accepted Manuscript version arising.

## Notes

### Competing Interest Statement

The authors have declared no competing interest.

### Summary of Updates

Accepted version. Only minor typographical changes

