## Supplementary material for "Analysis of coding gene expression from small RNA sequencing": supplentary information

**Supplementary Table 1**

| species | tissue | Sequencing depth | Sequencing depth (>25nt) | $R^2$ | Complexity |
|---|---|---|---|---|---|
| human | brain | 8303 | 3238 | 0.76 | 0.638 |
| human | cerebellum | 8098 | 1605 | 0.62 | 0.635 |
| human | heart | 8123 | 514 | 0.33 | 0.498 |
| human | kidney | 8644 | 1926 | 0.4 | 0.712 |
| mouse | brain | 11154 | 1196 | 0.52 | 0.712 |
| mouse | cerebellum | 11664 | 904 | 0.53 | 0.660 |
| mouse | heart | 14818 | 1130 | 0.69 | 0.707 |
| mouse | kidney | 9788 | 706 | 0.48 | 0.668 |
| mouse | testis | 11572 | 8518 | 0.41 | 0.780 |
| chicken | brain | 20838 | 448 | 0.56 | 0.747 |
| chicken | cerebellum | 19313 | 388 | 0.45 | 0.773 |
| chicken | heart | 28251 | 2696 | 0.71 | 0.756 |
| chicken | testis | 27806 | 15281 | 0.29 | 0.846 |

Sequencing depth is measured as the number of sequenced reads per megabase, considering the size of the reference genome used. $R^2$ is the coefficient of determination from the regression analysis comparing the expression levels of coding genes retrieved from small and total RNA sequencing libraries. Complexity refers to the Normalised Shannon entropy, defined as $\frac{\sum^{i} p_i \log_2 p_i}{\log_2(N)}$ where $p_i$ is the proportion of reads of type $i$ and $N$ is the total number of unique reads.

**Supplementary Table 2**

| | | Term (linear model) | | |
|---|---|---|---|---|
| **Tissue** | **Statistics** | **Total RNA** | **Half-life** | **Interaction** |
| **Brain** | **Slope** | 0.659 | -0.017 | -0.001 |
| | **SE** | 0.007 | 0.004 | 0.001 |
| | **p** | 0.000 | 0.000 | 0.452 |
| **Cerebellum** | **Slope** | 0.446 | -0.011 | 0.001 |
| | **SE** | 0.006 | 0.003 | 0.001 |
| | **p** | 0.000 | 0.001 | 0.098 |
| **Heart** | **Slope** | 0.246 | -0.018 | -0.001 |
| | **SE** | 0.006 | 0.003 | 0.001 |
| | **p** | 0.000 | 0.000 | 0.375 |
| **Kidney** | **Slope** | 0.273 | -0.019 | 0.002 |
| | **SE** | 0.007 | 0.004 | 0.001 |
| | **p** | 0.000 | 0.000 | 0.010 |

Statistical analysis of the linear model fitting the inferred gene expression levels from small RNA libraries against the levels from total RNA libraries and the half-life of the analysed transcripts.

**Supplementary Figure 1**

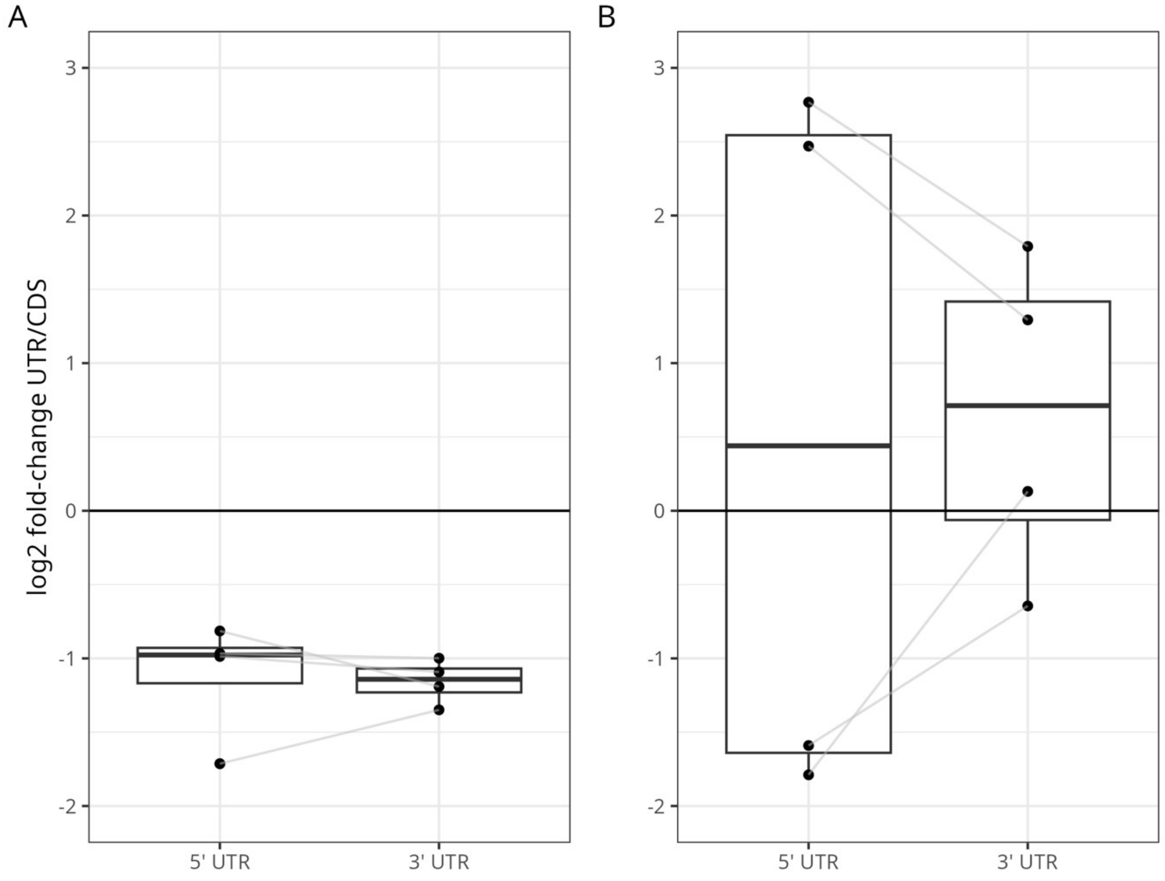


**Supplementary Figure 1. Relative mapping of sequencing reads to untranslated regions.** Log2 fold-change of the number of reads mapped to either the 5' untranslated region (5' UTR) or the 3' UTR with respect to reads mapped to the coding sequence. Values over 1 show an enrichment whilst value under 0 indicate a paucity of reads mapped to the UTR. Grey lines connect identical datasets (A) UTR fold-enrichment for reads from total RNA libraries. (B) UTR fold-enrichment for reads from small RNA libraries.

**Supplementary Figure 2**

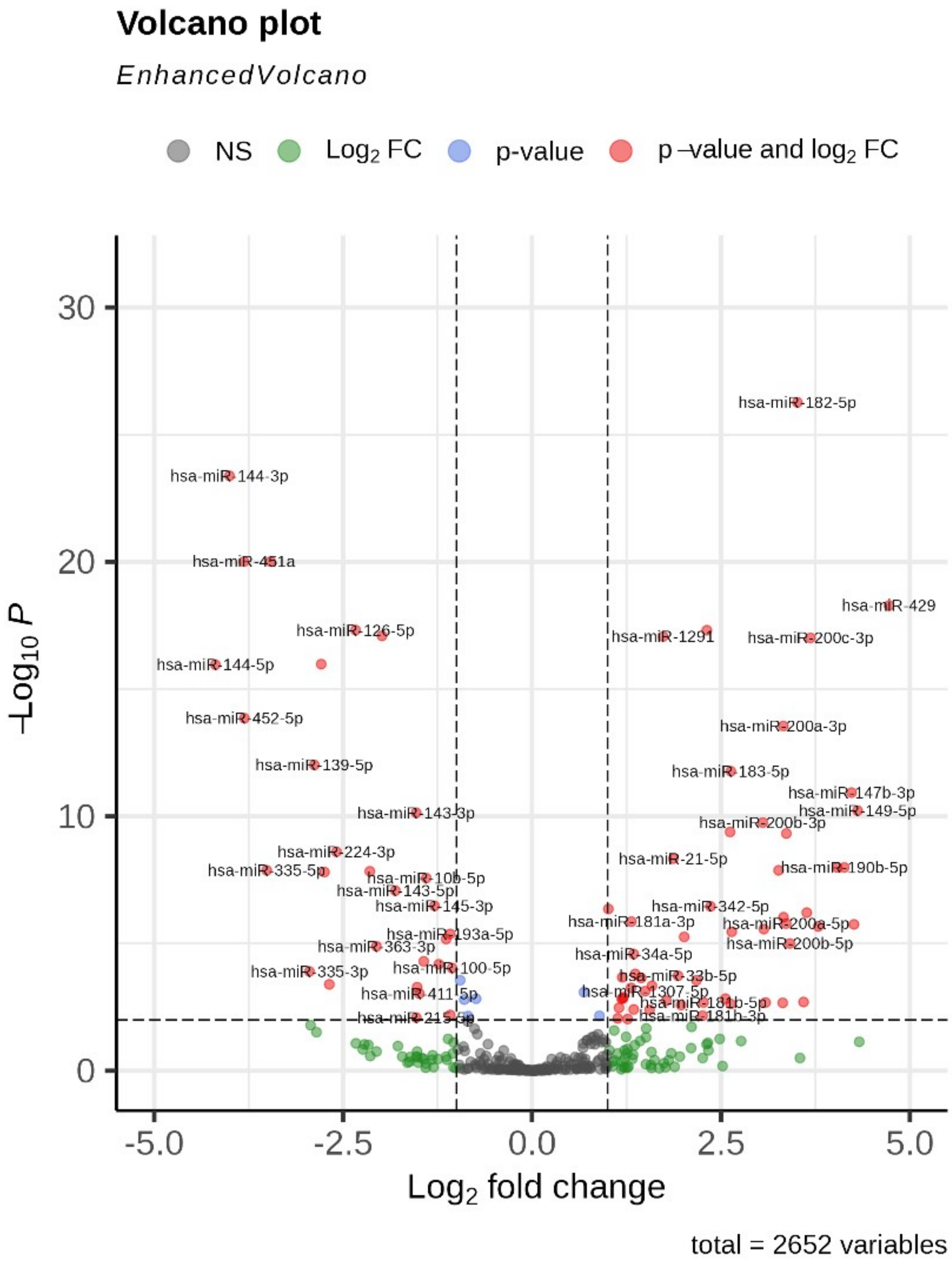


**Supplementary Figure 2. Differential gene expression of microRNAs from breast cancer paired sRNAseq experiments.** Volcano plot representing the expression fold change (DESeq2) in the x-axis of paired breast cancer samples (see main text) against the -log10 of the q value (FDR corrected p value) generated during the differential gene expression analysis. Identified differentially expressed genes are shown in red.

**Supplementary Figure 3**

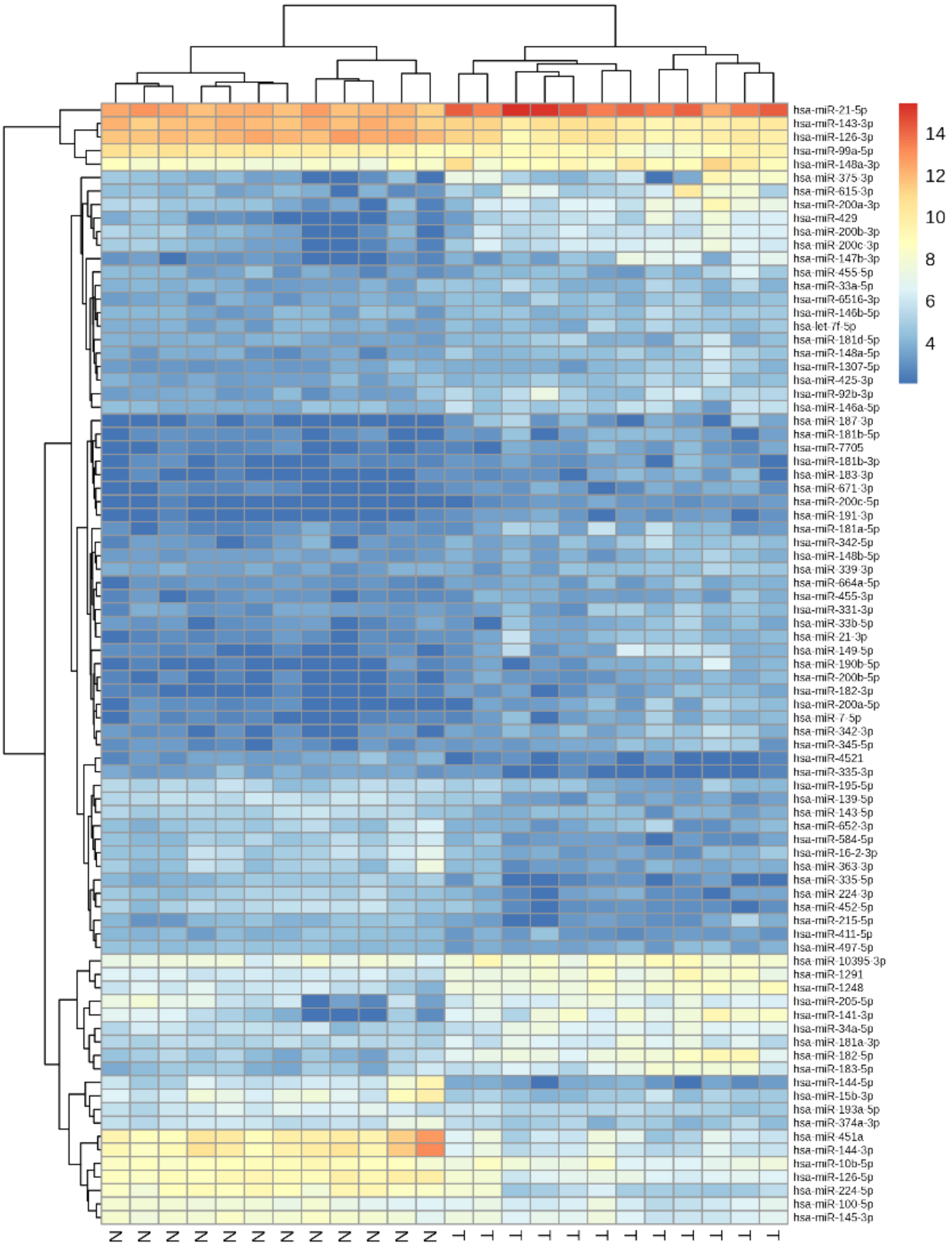


**Supplementary Figure 3. Gene expression profile of microRNAs from breast cancer.** Heatmap showing the expression levels of microRNAs from small RNA sequencing experiments in paired breast cancer samples.

**Supplementary Figure 4**

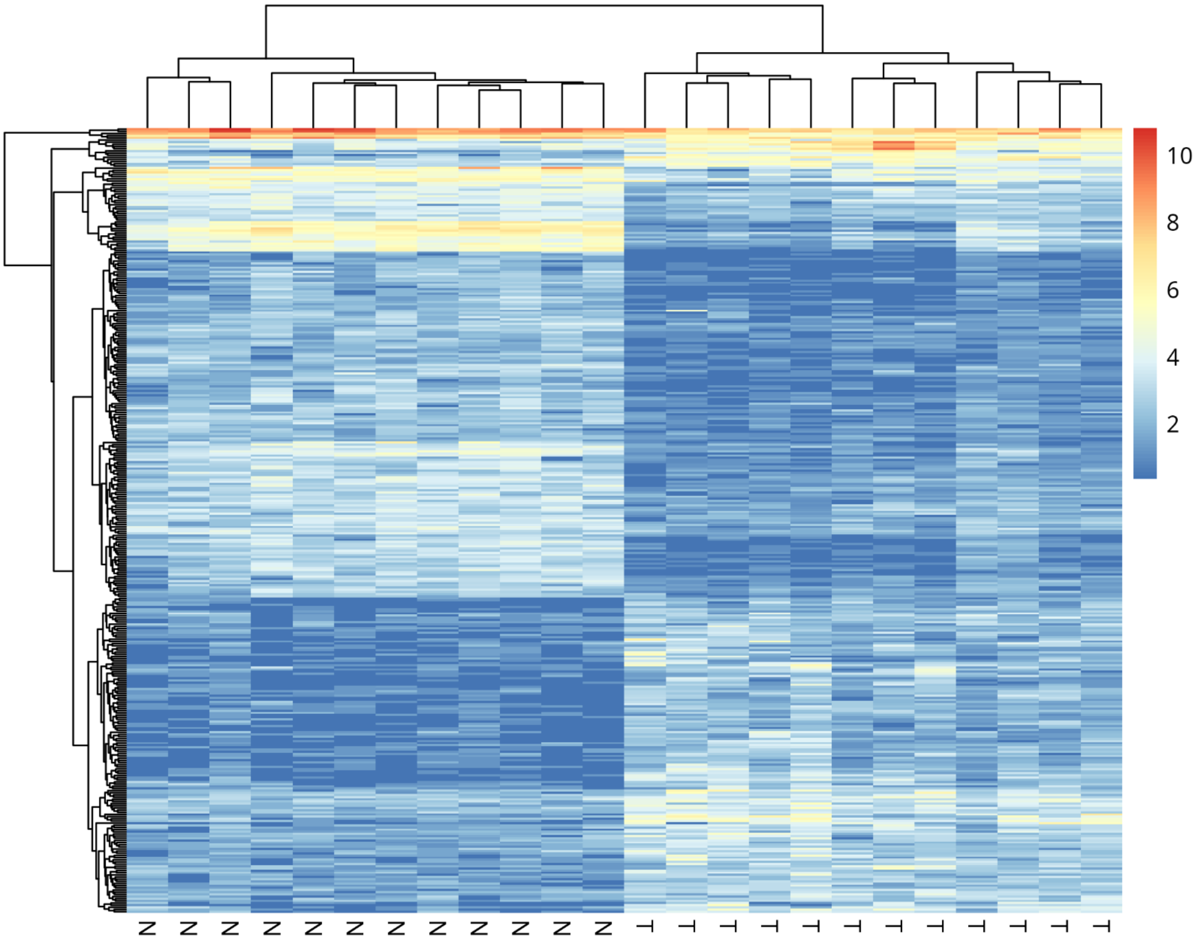


**Supplementary Figure 4. Gene expression profile of protein-coding transcripts from breast cancer.** Heatmap showing the expression levels of protein-coding genes from small RNA sequencing experiments in paired breast cancer samples.

**Supplementary Figure 5**

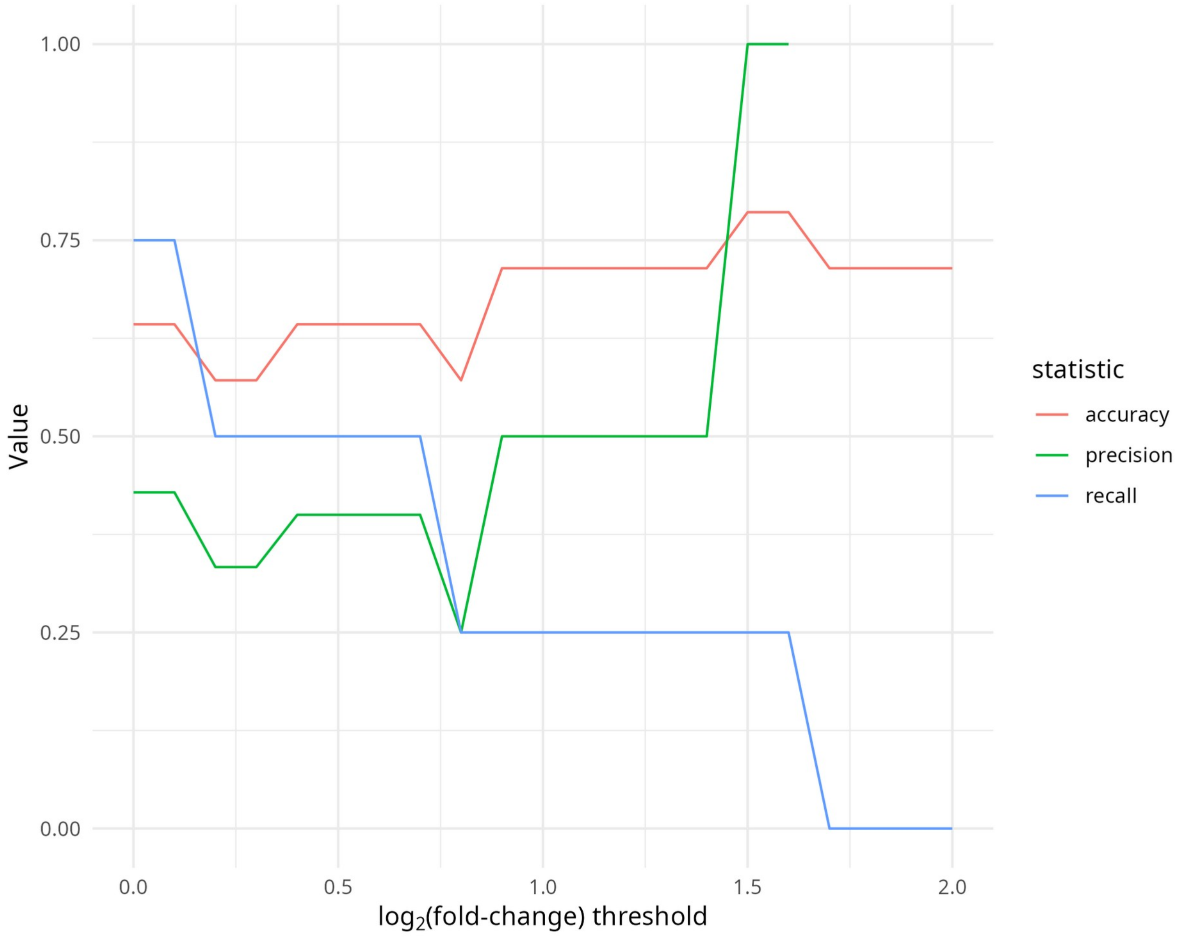


**Supplementary Figure 5. Accuracy, precision and recall when comparing the DGE analysis with the gold standard.** For thresholds of $\log_2$(fold-changes) in gene expression of the Differential Gene Expression analysis of Breast Cancer samples, the resulting upregulated genes were compared with the gold standard generated by Meerson et al. (2019). Values shown for accuracy, precision and recall.
